# Expression of glucocorticoid and mineralocorticoid receptor genes co-vary with a stress-related colour signal in barn owls

**DOI:** 10.1101/517060

**Authors:** Paul Béziers, Anne-Lyse Ducrest, Luis M. San-Jose, Céline Simon, Alexandre Roulin

## Abstract

Glucocorticoid hormones are important intermediates between an organism and its environment. They enable an organism to adjust its behavioural and physiological processes in response to environmental changes by binding to mineralocorticoid receptors (MR) and glucocorticoid receptors (GR) expressed in many tissues, including the integument. The regulation of glucocorticoids covaries with melanin-based colouration in numerous species, an association that might result from the melanocortin system and evolve within a signalling context. Most studies have focused on the circulating levels of glucocorticoids disregarding the receptors that mediate their action and could partly account for the covariation between the regulation of stress and melanin-based coloration. We investigated the association between melanin-based coloration and the gene expression levels of *GR* and *MR* but also of the melanocortin system and melanogenesis in the growing feathers of nestling barn owls (*Tyto alba*). We also considered the expression of genes related to the melanocortin system and melanogenesis to better understand the origin of the link between the expression of receptors to which corticosterone binds and melanin-based coloration. Nestling barn owls displaying larger black feather spots expressed *GR* and *MR* at lower levels than did smaller-spotted individuals. Because the expression of the *GR* and *MR* genes was positively, rather than negatively, correlated with the expression of melanogenic genes, we suggest that the link between melanin-based coloration, GR and MR is not directly associated to melanogenesis. The finding that the expression of *GR* and *MR* was positively associated with the expression of the *PCSK2* gene (encoding the protein convertase responsible for post-translational modification of the proopiomelanocortin hormone) suggests that the melanocortin system may be implicated in the establishment of the covariation between melanism and the expression of receptors to which glucocorticoids bind. Together with previous studies, our results suggest that coloration is not only related to the production of glucocorticoids but also to the sensitivity of the integument to glucocorticoids.

## Introduction

Hormones play an important role in translating external stimuli into physiological changes to enhance the survival prospects of an organism in the face of continuous changes in its environment [53]. The glucocorticoid hormones: corticosterone in amphibians, reptiles, birds and rodents and cortisol in non-rodent mammals and fishes, are under the control of the hypothalamic-pituitary-adrenal (HPA) axis and affect most cells in the body [6, 61]. Thus, variation in the levels of glucocorticoid hormones (hereafter referred to as “corticosterone”) plays a major role in many physiological and behavioural functions [reviewed in 98, 113], having important consequences on the development [66, 109], health [22, 41], reproductive success [11, 15, 108] and survival of an individual [39, 82, 84]. Such multiple links to aspects of an individual’s fitness have been proposed to favour the evolution of animal ornaments that reflect corticosterone levels or the capacity of an individual to withstand stressful conditions [16, 45]. However, although several empirical cases support the expected association between corticosterone levels and different types of ornaments (coloration and other visual signals, vocalizations, among others [see also 17, 64, 67, 88, 96, 111]), the proximate factors driving it remain obscure for most types of ornament, particularly for those whose expression shows little environmental plasticity, such as colorations based on the deposition of melanin pigments [87].

In many vertebrate species, corticosterone is associated with melanin-based colorations [97]. Most studies that have investigated the covariation between melanin-based colour traits and the HPA axis have focused on glucocorticoid levels in plasma [e.g., 21, 54, 92] or in integuments, such as feathers and hairs [9, 37, 52]. Although the HPA axis is a complex system regulated at multiple levels and by multiple factors [13, 34, 44, 83], it is clear that the action of corticosterone is largely mediated in a dose-dependent manner by the amounts of glucocorticoid (GR) and mineralocorticoid (MR) receptors that bind to corticosterone in different target tissues [46, 106, 114]. The nuclear mineralocorticoid (MR) and glucocorticoid (GR) receptors are ligand-dependent transcription factors that influence the expression of numerous genes implicated in growth, metabolism, reproduction, resource allocation and immune functions [23, 63, 81]. The nuclear MR has a ten-fold higher binding affinity to glucocorticoids than GR which is, however, more numerous and hence have a higher carrying capacity than MR [24, 33]. These differences in properties are thought to mediate independently the action of baseline (MR) and stress-induced glucocorticoid levels (GR). Thus, MR is mainly thought to regulate the circadian variation of glucocorticoid levels while GR is mainly bound after an acute challenge or at the peak of ultradian rhythm when glucocorticoid concentration is high [20], implying that MR are rapidly saturated while unbound GR are still numerous.

Glucocorticoid receptors are well distributed throughout the body with GR being more ubiquitous than MR [6, 34, 61, 62]. Their distribution can show seasonal variations across tissues [12, 60, 62], and be influenced or not by circulating hormone levels or stress [14, 26, 60]. More importantly, glucocorticoid receptors have been found to be expressed in different skin cells like melanocytes where melanin pigments are produced [100]. Measuring variation in the expression of glucocorticoid receptors (GR, MR) is thus key to not only understand how the activity and action of the HPA axis is regulated, but also how corticosterone levels influence the development of melanin-based colorations and why corticosterone levels covary with melanin-based colorations. Despite their importance in the regulation of the HPA axis and despite that they have been shown to relate to fitness (*e.g.*, receptors can vary in certain tissues between individuals adopting different reproductive tactics: Artbery, Deitcher & Bass 2010), only one study has investigated the link between melanin-based traits and the amount of glucocorticoid (GR) and mineralocorticoid (MR) receptors [59]. In this study, conducted in house sparrows (*Passer domesticus*), males with large black bibs showed a fewer amount of GR and MR in skin tissues than males with small bibs but the generality of this finding remains untested.

Additionally, it is still unclear by which means glucocorticoid (GR) and mineralocorticoid (MR) receptors and, thereby corticosterone, could covary the synthesis of melanin in the integument. The association between corticosterone and melanin-based coloration has been proposed to originate from the pleiotropic effect of the melanocortin system [30, 89] or alternatively, because adrenal glands and pigment cells (melanocytes) both originate from the neural crest cells [112]. The melanocortin system is composed of five transmembrane melanocortin receptors that bind different melanocortin hormones and regulate tissue specific processes, including melanin synthesis and stress response. On the one hand, in hair and feather follicles, binding of the melanocortin α-MSH (α-melanocyte-stimulating hormone) to the melanocortin 1-receptor (MC1R), induces the synthesis of melanin pigments. On the other hand, binding of the melanocortin ACTH (adrenocorticotrophin hormone) to the melanocortin 2-receptor (MC2R) expressed in the adrenal glands and the skin activates the downstream signalling for the production of corticosterone [51, 101]. Both melanocortin hormones, α-MSH and ACTH, are derived from the cleavage of the proopiomelanocortin (POMC) prohormone by the protein convertases PC 1 and PC 2 [19, 31, 78]. Thus, the link between melanin-based coloration and the HPA axis could potentially arise because α-MSH and ACTH are derived from the same precursors and both modulated by the same convertases in the same tissue. However, additional empirical knowledge is still needed to confirm this hypothesis.

In the present study, we examined the relation between melanin-based traits and the expression levels of *GR* and *MR* in growing feathers of nestling barn owls on the breast and belly, which can present very different colour patterns. The barn owl (*Tyto alba*) displays melanin-based traits that are associated to various life-history, morphological, behavioural and physiological traits [86]. Barn owls displaying larger black spots on the feather’s tip of their ventral body side are less sensitive to stress than smaller-spotted individuals, as they have a lower increase in blood corticosterone levels after a stressful event [1], are less affected by an artificial increase in blood corticosterone levels during growth and breeding [2, 3] and loose less weight under food depletion [27]. Thus, given that large spotted individuals are thought to be less stress-sensitive than small spotted individuals, we predict that large spotted individuals should express less *GR* and *MR* in their feathers than small spotted individuals. In parallel, we measured the expression levels of different melanocortin- and melanogenesis-related genes in order to identify the potential proximate mechanism or mechanisms linking melanin-based colour traits and the HPA axis activity. Assuming that genes involved in melanogenesis, or those of the melanocortin system, are involved in the association between melanin-based coloration and the expression levels of *GR* and *MR*, we expect to find a negative relation between the expression of melanin and melanocortin-related genes with receptors involved in the binding of glucocorticoids (*GR, MR*), given the previously described negative relation between corticosterone and melanin-based traits in barn owls.

## Methods

### Study species and site

The barn owl is a medium sized raptor that lives in open landscapes and preys on small mammals. Females lay two to eleven eggs at a rate of one egg every two to three days between February and the beginning of August. After an incubation of ca. 32 days, nestlings hatch asynchronously every two to three days, which leads to a within-brood age hierarchy [85]. Females stay in the nest box until the elder nestlings are about 3 weeks old, able to thermoregulate by themselves and to feed without maternal help. Pennaceous feathers start growing around 10-15 days of age and plumage is completed at approximately 55 days, when nestlings are ready to fledge.

The study was carried out in 2014 in an open rural landscape in western Switzerland (46°49’N, 06°56’E) and based on a sampling of 48 male and 35 female nestlings from 26 breeding pairs. Nestlings were visited for the first time at an early stage of feather development, when they were between 15 and 30 days old (mean ± SD: 21.81 ± 3.7 days). During this visit, we ringed the nestlings, plucked 3 to 5 feathers from their breast and belly, and took a blood sample for molecular sex determination following the protocol described by Py et *al*. (2006) [79]. We plucked feathers at an early stage of development to pinpoint the time when nestlings produce the black eumelanic spots on the upper part of the feathers. In this study, feathers were plucked at an earlier age compared to the previous study of San-Jose et *al.* (2017) [95] where spots were already formed. We re-visited the nest-boxes a second time when nestling feathers were fully developed (at ca. 55 days of age) in order to assess plumage coloration (see below for procedure). At each visit, we also weighed the nestlings and measured the length of their left wing in order to precisely determine their age. Most of the nestlings (75 out of 83) were cross-fostered before hatching between randomly chosen nests to allocate genotypes randomly among rearing environments and to avoid the potential confounding effect of the biological mother on the relationship between nestling spot size and the expression levels of *GR, MR*, melanogenic genes and genes belonging to the melanocortin system. For this, we cross-fostered eggs between pairs of clutches showing similar laying dates. We swapped all eggs between pairs of clutches that were of similar size (± 1 egg), whereas if there was a difference of more than one egg, we cross-fostered the same number of eggs between pairs of randomly chosen nests. In this case, we made sure to keep intact the within-brood age hierarchy by swapping eggs at the same stage of incubation. We captured adult females and measured their plumage coloration while they were incubating eggs or raising their chicks.

### Assessment of plumage coloration

In the Swiss population of barn owls, the overall colour of ventral body side varies from white to rufous owing to differential deposition of pheomelanin, whereas the tips of the ventral feathers are covered with a varying number of black spots, whose diameter and number also varies between individuals. On average, females are redder and display larger and more eumelanic spots than males [28]. Variation in the pheomelanin-based coloration is in part due to a valine (V)-to-isoleucine (I) substitution at the amino acid position 126 of the melanocortin 1-receptor (MC1R) [94], which is an important component of the melanin synthesis. This mutation was also shown to differently affect the expression of melanogenic-related genes [95]. Individuals presenting the *MC1R* white allele (*MC1R*_*White*_) express the *MC1R* and melanic genes at lower levels (*TYR, TYRP1, OCA2, SLC45A2, KIT, DCT*) than individuals carrying the *MC1R* rufous allele (*MC1R*_*Rufous*_) responsible for the rufous coloration. The expression of these plumage traits is strongly heritable and weakly sensitive to the rearing environment [90]. In the present study, we focused only on the size of the black spots given that in previous studies no associations have been found between spot number or the pheomelanin-based coloration with stress-related parameters [1, 2]. Within a 60 x 40 mm2 frame placed on a standard position of the breast and the belly of all nestlings and adults we measured the diameter of 10 to 15 black spots to the nearest 0.1 mm using a slide calliper. We used the mean diameter of spots per body part for our statistical analyses. These methods have already been shown to be reliable [86].

### RNA extraction

We extracted total RNA from the base of the developing feathers plucked from the breast and belly of the nestlings. We used breast and belly feathers because they can present very different colour patterns in terms of spots size. Additionally, by using feathers from two different body parts, we can assess whether an association between GR, MR and melanin-based traits is related to melanogenic genes (*MLANA, MITF-M* a melanocyte-specific isoform of *MITF, OCA2, PMEL, TYR)* and to genes of the melanocortin system (*PCSK 2, MC1R*) in a consistent manner between body parts. The feathers were collected within the hour following the capture of the nestlings and directly stored in liquid nitrogen before being stored at −80°C until gene expression analyses. The sampling time (*i.e.*, the time between capture and blood sampling the nestling) was not considered in our models given that it had no effect on the level of expression of the different genes (not shown).

For each nestling and body part, two to five feather bases were grounded with a pestle in liquid nitrogen, resuspended in 600 µL of lysis buffer containing 10 mM DTT and passed through a spin filter column (InviTrap Spin Universal RNA Mini kit, Stratec, Berlin, Germany) to remove genomic DNA, barbs and barbules. Following the protocols (InviTrap Spin Universal RNA Mini kit) total RNAs were eluted in 50 µL of water. To further remove all possible genomic DNA contaminants, a second DNAse I treatment was applied by incubating the 50 µL of eluted RNA with 40 U of DNAse I (Roche diagnostics Ltd, Basel, Switzerland), 50 U of RNase Inhibitor (Promega AG, Dübendorf, Switzerland) and 1x Roche buffer for 30 min at 37°C, before rapidly freezing the solution at −80°C. The RNA was then purified and concentrated to 20 µL with the GeneJet RNA purification Kit (Thermo Fisher Scientific, Ecublens, Switzerland). For each series of RNA extraction (12 samples), one random sample was analysed with the Fragment analyser of Advanced Analytical (Labgene, Châtel-St-Denis, Switzerland) to assess RNA quality. All RQN (RNA quality number) values were above 8.0 (scale from 1 to 10), which indicates that the RNA samples were of high quality [68, 115]. Finally, 1 µg of DNase I-treated total RNA was reverse transcribed in a final volume of 20 µL, using 4 µL of 5x VILO reaction buffer and 2 µL of 10x SuperScript Enzyme Mix reverse transcriptase (Thermo Fisher Scientific) at 25°C for 10 min, at 42°C for 60 min followed by reverse transcriptase inactivation at 85°C for 5 minutes. The cDNA was diluted 10x with 1x TE (10 mM Tris-HCl, pH 8.0, 0.1 mM EDTA) to achieve a final quantity of 100 ng of RNA and finally stored at **-**20°C until quantification.

### Gene sequencing

To explore the potential proximate mechanism linking melanin-based traits and the HPA axis activity, we choose to determine the relationship between genes involved in the HPA axis (*GR, MR*) with genes involved in the melanocortin system (*PCSK 2, MC1R*) and the synthesis and development of melanin pigments (*MLANA, MITF-M* a melanocyte-specific isoform of *MITF, OCA2, PMEL, TYR)*. We did not measure the expression of the *POMC* gene, a main actor of the melanocortin system, because in barn owl feathers we only detect its truncated form, which may not be functional [95]. The convertase PC2 encoded by the gene *PCSK 2* cleaves the POMC prohormone into different melanocortin peptides (*e.g.*, ACTH, α-MSH) that bind to and activates different melanocortin receptors, including MC1R and MC2R [19, 31, 35, 78]. Binding of the ligand α-MSH to the MC1R receptor triggers the melanogenesis process by activating the expression of the microphthalmia transcription factor (*MITF*) gene. MITF is a transcription factor that upregulates the expression of different genes involved in the synthesis of melanins (*e.g.*, tyrosinase, *TYR*) [29, 105] and in the development and maturation of the melanosomes (*e.g.*, protein melan-A (*MLANA*), Pre-MELanosome protein (*PMEL*), melanocyte-specific transporter protein (*OCA2*)). MLANA is a transmembrane protein expressed on the surface of melanosomes and plays an essential role in melanosome biogenesis, vesicular trafficking and melanosome maturation and autophagy [5, 38, 43]. The *PMEL* gene codes for a transmembrane glycoprotein that is regulated by multiple proteolytic processes resulting in an amyloid fibrillar matrix, which enables the deposition of melanin pigments and potentially prevents the toxic effect of melanin derivatives in the melanosome [10]. The *OCA2* gene, also known as the pink-eyed dilution protein or p-gene in human, plays an important role in the biogenesis of melanosomes and regulation of eumelanin content in melanocytes through the processing and trafficking of the tyrosinase enzyme [42, 71, 72, 104].

The genes *GR* (*NR3C1), MR* (*NR3C2), MLANA, MITF-M, PMEL* and *TBP* were sequenced from polyA cDNAs following the protocols described in San-Jose et *al.* (2017) [95]. We sequenced the 8 exons of the glucocorticoid and mineralocorticoid receptors (*GR* and *MR*). The 2380 bp of *GR* included the start codon (position 1) and the stop codon (2322 bp). We found two synonymous mutations c.582C>A (I194, with 6 individuals of genotype *CC*, 1 of genotype *AA* and 1 heterozygous individual *CA*) and c.1557G>A (A519, with 6 homozygous individuals *GG*, 1 *AA* and 1 heterozygous *GA*) (for accession numbers see Table S1). The 3108 bp sequence of the *MR* gene contained the start and stop codons located at 112 bp and 3055 bp, downstream the beginning of the sequence, respectively. We found four mutations, three of them were synonymous: c.261C>T (C87, 3 *CC*, 2 *TT* and 3 *CT* individuals), c.1353T>C (S451, 8 *TT*, 0 *CC* and 2 *TC*) and c.1707G>T (T569, 7 *GG*, 0 *TT*, 3 *TG*), and one non-synonymous: 292C>G (Q98E, 3 *CC*, 1 *GG* and 4 *CG*). We sequenced 590 bp corresponding to the 5 exons of *MLANA* (*MART-1*) with a start and stop codon at 69 and 410 bp downstream of the begin of the sequence, respectively. One non-synonymous mutation at position c.237C>T resulted in an arginine to threonine substitution (R56T, with 11 homozygotes *CC* and 1 heterozygous *CT* individuals). In addition, we sequenced ten exons corresponding to 1977 bp of the Pre-MELanosome protein gene (*PMEL*), also called *gp100* or *SILV*. We also sequenced 1483 bp of the TATA-binding protein (*TBP*) and 1054 bp of the glyceraldehyde-3-phosphate deshydrogenase (*GAPDH*) which were used with the ribosome protein L13 (*RPL13*) sequenced in a previous study [95], as reference genes for qPCR. The sequence of TBP contained the start and the stop codons at position 135 and 1043 bp and includes 7 exons and the GAPDH sequence contains at least 10 exons but was missing the start codon. None of the mutations were related to barn owl coloration or to gene expression. Sequences of all the other genes (*MC1R, OCA2, PCSK 2, RPL13, TYR*) are described in a previous study [for details see 95].

### Quantitative PCR

The specific qPCR primers and dual labelled fluorescent probes (FAM-BHQ1, fluorescein with BHQ1 quencher) were designed with Primer 3 (http://bioinfo.ut.ee/primer3-0.4.0/primer3/) and DNASTAR PrimerSelect software or directly by Microsynth (Balgach, Switzerland). Each primer and probe was blasted to verify its specificity before purchasing them from Microsynth (Balgach, Switzerland) and Eurogentec (Liège, Belgium). We then assessed the concentrations for each gene primer pairs and probes in order to reach a qPCR amplification efficiency of 95 to 105% (for details on efficiency see Table S2 in the supplementary material). Due to the low levels of gene expression of some genes (*MC1R, PCSK2*), we pre-amplified the cDNA with the TaqMan preamplification Master Mix kit (Thermo Fisher Scientific) for 14 cycles. We verified that the pre-amplification was uniform by calculating the delta**–**delta Ct values, which are ΔCt of the pre-amplified gene minus ΔCt of the cDNA (Ct values of the target genes minus Ct values of the reference gene *GAPDH*). The qPCR assays were run on a QuantStudio 6 real-time PCR system (Thermo Fisher Scientific) and each sample was run in triplicate with 2 µL of the diluted cDNA and 8 µL of master mix containing 1x qPCR MasterMix Plus Low ROX (Eurogentec, Liège, Belgium) or 1x TaqMan gene expression MasterMix (Thermo Fisher Scientific) and primers and probes (for more details on primers and probes concentration see Table S2). To correct for interplate variation, each pre-amplified cDNA sample was arbitrarily assigned to a plate and three pools of different pre-amplified cDNAs were added to each plate and used to set up the threshold values between plates. To correct for any variation in cDNA content, CT scores of the candidate genes were normalized using three reference genes: the ribosome protein L13 (*RPL13*), glyceraldehyde-3-phosphate deshydrogenase (*GAPDH*) and TATA-binding protein (*TBP*) (for accession numbers see Table S1). CT scores were imported into qBasePLUS software 1.3 (Biogazelle, Zwijnaarde, Belgium) and reference genes quality analysed with geNorm [107]. We calculated the mean relative quantities (RQs) per sample and used these values for our statistical analyses. All the genes were transformed to fulfil the normality assumptions of parametric statistical tests.

### Statistical analyses

We used linear mixed effect models to test if the expression of mineralocorticoid (*MR*) and glucocorticoid (*GR*) receptor genes (*MR* and *GR* were considered as dependent variables in separate models) is related to the size of eumelanic spots displayed on the tip of ventral feathers and/or the expression of genes associated to the melanocortin system (*MC1R, PCSK 2*) and melanogenesis (*MLANA, MITF-M, OCA2, PMEL, TYR*). To obtain a reduced set of genes associated to the melanocortin system and to the deposition of melanin pigments, we first performed a principal components (PC) analysis and limited the number of components to the number that accounted for 80% of the total variance [50, Table 1]. The PCA yielded 3 components that explained 81.3% of the variance. The first PC represents genes associated to the melanogenesis pathway (*PMEL, MLANA, TYR*) with a gene associated to the melanocortin system and the melanic process (*MC1R*). The 2nd PC is mostly represented by one melanogenic gene (*MITF-M*). Whereas the 3rd PC represents a gene (*PCSK 2*) involved in the melanocortin system.

**Table 1:** Principal components analysis of eight genes associated to the melanocortin system (*MC1R, PCSK2*) and melanogenesis (*MLANA, MITF-M, OCA2, PMEL, TYR*). Correlation between the original variables with their standard errors and the three first components (PC 1, PC 2, PC 3), eigenvalues and cumulative variance in percentage. The coordinates of individuals of first three components were used in the mixed effect models. Values in bold and *italic* represent parameters with, respectively, a high and moderate correlation (0.4 to 0.6) with components.

| Genes | PC1 | PC2 | PC3 |
| --- | --- | --- | --- |
| *PCSK 2* | 0.26 ± 0.05 | 0.31 ± 0.03 | **-0.79 ± 0.02** |
| *TYR* | -0.42 ± 0.09 | 0.23 ± 0.06 | -0.05 ± 0.09 |
| *MC1R* | -0.43 ± 0.12 | -0.26 ± 0.06 | -0.27 ± 0.07 |
| *PMEL* | -0.49 ± 0.12 | -0.14 ± 0.08 | -0.22 ± 0.08 |
| *MLANA* | -0.47 ± 0.14 | -0.04 ± 0.17 | -0.22 ± 0.15 |
| *OCA2* | -0.33 ± 0.16 | *0.47 ± 0.12* | *0.45 ± 0.08* |
| *MITF-M* | -0.04 ± 0.08 | **0.74 ± 0.07** | -0.08 ± 0.08 |
| Eigen values | 3.4 | 1.4 | 0.9 |
| Cumulative variance | 48.5 | 68.6 | 81.3 |

In the models, we considered as fixed factors the mean spot diameter of the nestlings, the body part where the feathers were plucked from, *i.e.*, the breast and belly, and the three PCs of gene expression (PC 1, PC 2, PC 3, Table 1). The date (*i.e.*, Julian date) and time of the day (hour) when the feathers were sampled as well as sampling time (*i.e.*, time between the moment when a nestling was captured and its feathers plucked), sex, body mass and age of the nestlings were not considered in the models as preliminary analyses showed that these variables were not associated to the expression of *GR* and *MR* receptors (for each of these variables the 95% credible interval (CrI) overlapped with zero). Previous analyses also showed no interaction between body parts and the three PCs and spot diameter and were therefore removed from the models (for each of the interactions the 95% credible interval (CrI) overlapped with zero). Spot diameter was not correlated to any of the PCs (all 95% confidence intervals overlapped with zero) and was thus considered in the same model. The random effect of brood identity was considered in the models as well as the random effect of nestling identity in order to consider the repeated measurements taken on the nestlings (*i.e.*, the observed values at the two different body parts), with the random effect identity of nestling nested in the brood of origin.

All statistical analyses were done with R 3.5.1 [80]. Linear mixed effect models were run with *lme4* package [8] and principal components with the *prcomp* function from the *stats* package. The significance of predictors were assessed by simulating 50’000 random values from the joint posterior distribution of the model parameters using the *sim* function from the *arm* package [36]. We used the 2.5 % and 97.5 % quantiles of the simulated data as the lower and upper bands of our 95% CrI [55]. Predictors with credible interval that not include zero were considered to be significantly associated to the expression levels of GR and MR receptor genes. To compare effect sizes of individual model terms, all numerical variables were standardized to mean of zero and SD of one. For each of the models, we visually inspected the residuals to control that assumptions of normality and heterocedasticity for mixed effect models.

For the figures, we simulated 5’000 random samples from the joint posterior distribution of the model parameters within the all range of the predictor variables (or each factor level). From these simulations, we used the mean predicted values and the 2.5 % and 97.5 % quantiles as lower and upper limits of the 95 % CrI.

## Results

### Co-expression of glucocorticoid and mineralocorticoid receptor genes with melanocortin and melanogenesis-related genes

The glucocorticoid receptor (*GR*) gene was positively associated to PC 2 and negatively to PC 3 which was the variable with the largest effect size (Table 2, Figure 1). A high factor score for PC 2 represents a high level of expression for the melanic genes *MITF-M, OCA2* and to a lower extent of the melanocortin gene *PCSK2* and a low expression level of the *MC1R* receptor gene. A high factor score for PC 3 represents a low level of expression of the *PCSK2* gene and a high expression for the melanic gene *OCA2*. In sum, the expression of the *GR* gene was positively co-expressed with *PCSK2, MITF-M* and *OCA2* genes and negatively co-expressed with *MC1R* receptor gene. Individuals with small spots expressed *GR* at higher levels than conspecifics with larger eumelanic spots (Table 2, Figure 3a). On the other hand, expression levels of *GR* were not differently expressed between body parts and not associated to PC 1.

**Table 2:** Relation between expression level of glucocorticoid (*GR*) and mineralocorticoid (*MR*) receptors and genes associated to the melanocortin system and melanogenesis pathway. Linear mixed effect model with brood identity and nestling identity nested in brood identity were introduced as random factors. Analyses were based on 156 feather samples taken on breast and belly of 81 nestling barn owls from 25 broods. Values in *italic* represent significant parameters.

| Predictors | *GR model estimates*<sup>†</sup><br>(95% CrI) | *MR model estimates*<sup>†</sup><br>(95% CrI) |
| --- | --- | --- |
| Intercept | 0.25 (0.17; 0.34) | 0.17 (0.1; 0.25) |
| PC1 | -0.02 (-0.05; 0.01) | *-0.06 (-0.09; -0.03)* |
| PC2 | *0.07 (0.02; 0.11)* | *0.15 (0.11; 0.2)* |
| PC3 | *-0.14 (-0.2; -0.08)* | *-0.09 (-0.15; -0.04)* |
| Size of eumelanic spots | *-0.08 (-0.14; -0.02)* | *-0.08 (-0.13; -0.02)* |
| breast - feather * | *-0.07 (-0.2; 0.06)* | *-0.05 (-0.16; 0.07)* |

<sup>†</sup> estimates are based on standardized predictors
\* gene expression difference from breast to belly feathers

**Figure 1:**
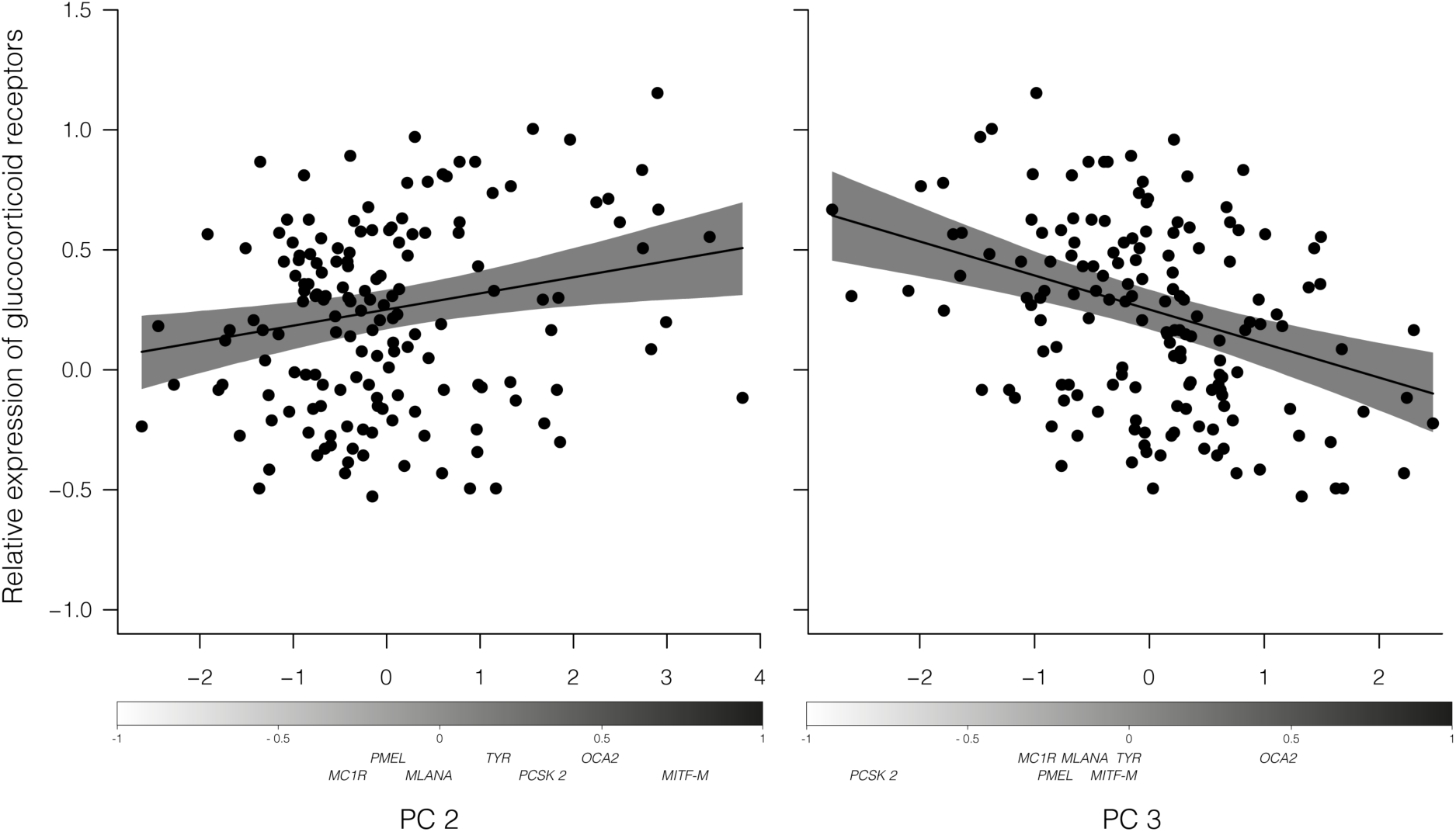
Relation between principal components (PC 2, PC 3) and the log-transformed expression of glucocorticoid (*GR*) receptor genes in feathers of nestling barn owls. The dots represent the raw data and the black lines represent the fitted values estimated from the GR model (Table 2). The grey shadings represent the 95% CrI of the regression lines. The gradient bars below the X axis represent the loadings (correlations) of each variable on each principal component (see Table 1).

The mineralocorticoid receptor (*MR*) gene was negatively associated to PC 1 and PC 3 and positively associated to PC 2 (Table 2, Figure 2). A high factor score for PC 1 represents a low level of expression of genes involved in melanin deposition (*TYR, PMEL, MLANA*) and receptor *MC1R*. Thus, the expression of *MR* receptor gene was positively co-expressed with melanic genes (*TYR, PMEL, MLANA, OCA2, MITF-M*) and negatively with *MC1R* receptor gene. Individuals with small spots expressed *MR* receptor gene at higher levels than conspecifics with larger eumelanic spots (Figure 3b). Expression of *MR* receptor gene was not differently expressed between body parts.

**Figure 2:**
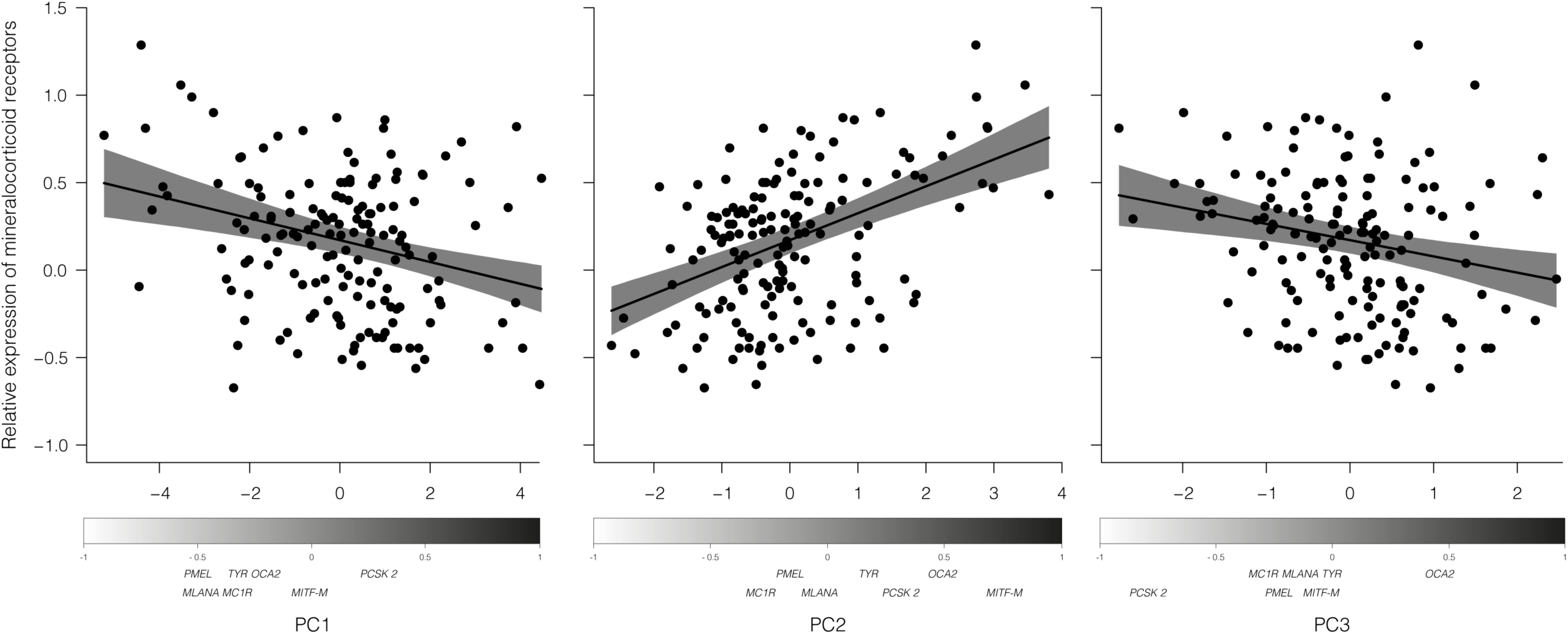
Relation between principal components (PC 1, PC 2, PC 3) and the log-transformed expression of mineralocorticoid (*MR*) receptor genes in feathers of nestling barn owls. The dots represent the raw data and the black lines represent the fitted values estimated from the MR model (Table 2). The grey shadings represent the 95% CrI of the regression lines. The gradient bars below the X axis represent the loadings (correlations) of each variable on each principal component (see Table 1).

**Figure 3:**
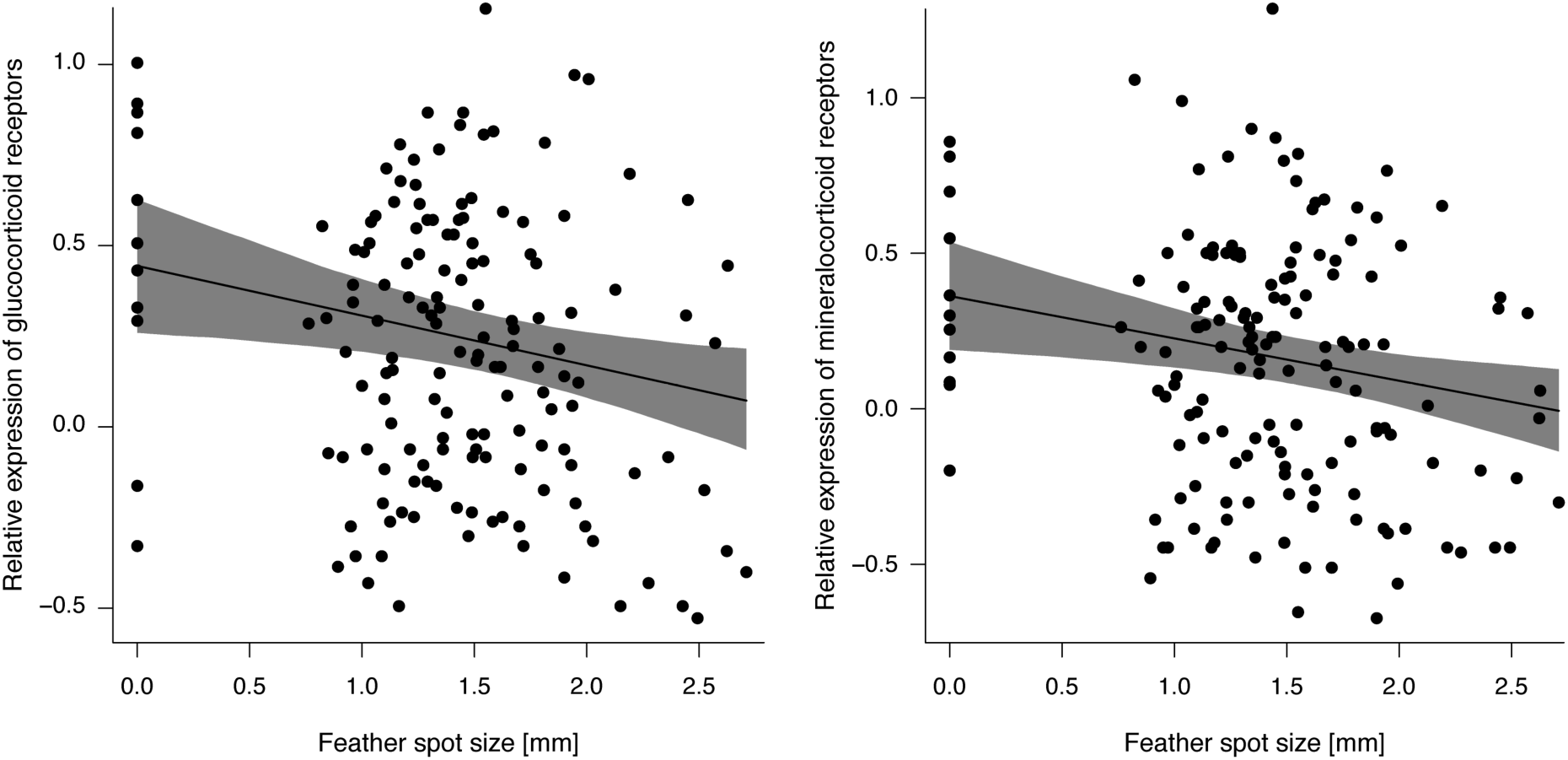
Relation between the size of black feather spots and the log-transformed expression of (a) mineralocorticoid (*MR*) and (b) glucocorticoid (*GR*) receptor genes in feathers of nestling barn owls. The dots represent the raw data and the black lines represent the fitted values estimated from MR and GR model (see Table 2). The grey shadings represent the 95% CrI of the regression lines.

## Discussion

Nestling barn owls displaying large eumelanic spots showed lower expression of glucocorticoid (*GR*) and mineralocorticoid receptors (*MR*) genes in breast and belly feathers compared to small-spotted conspecifics. These associations are in line with previous findings showing that the size of black feather spots is negatively related with the regulation of corticosterone, the hormone that binds to GR and MR in the barn owl [1, 2] and in other species [review in 30, 89]. Our results are also consistent with a previous study in house sparrows showing that males displaying large melanic bibs show less GR proteins in the skin of their bib compared to males with small bibs [59].

The relationship between melanin-based traits and the expression of glucocorticoid receptors (*MR, GR*) could arise for different reasons. One possibility is that this link arises through the melanocortin system and its multiple effects on the expression of melanogenic genes and the HPA axis. For instance, through the expression of the POMC prohormone or protein convertases (PCSK 1/3, PCSK 2) involved in the cleavage of the POMC prohormone into different active peptides [77]. The relationship could arise because the proteins involved in the processing of POMC, or the downstream effector products regulated by the melanocortin system, affect simultaneously the expression of GR and MR and melanin-based traits. The cleavage of the POMC prohormone results in the adrenocorticotrophic hormone (ACTH) that induces the production of corticosterone in adrenal glands and in skin [31, 102, 103], which is itself known to affect the expression of glucocorticoid receptors [49, 69, 74] and therefore, influence the abundance of MR and GR in skin tissues. ACTH can be further cleaved by the protein convertase 2 (PCSK 2) into corticotropin-like intermediate peptide (CLIP) and α-melanocyte-stimulating hormone (α-MSH), the latter being involved in the synthesis of melanin pigments once bound to the melanocortin 1-receptor (MC1R). The relationships between PC 2 and PC 3 with *GR* and *MR* could suggest that the melanocortin system and the upstream regulators PCSK 2 are implicated in the link with melanin coloration owing to its role in the cleavage of ACTH into α-MSH and CLIP. This relationship might be the result of a trade-off between the production of ACTH and α-MSH by PCSK 2. We can hypothesize that individuals expressing *PCSK 2* at a higher level may produce less ACTH and therefore less corticosterone but more receptors (*GR, MR*) and α-MSH and therefore be more melanic, whereas individuals with low expression levels of *PCSK 2* might produce more ACTH but less α-MSH. Even though we did not measure the corticosterone levels of nestlings in this study, this hypothesis is supported by previous studies in barn owls showing that melanin-based traits are negatively associated to circulating corticosterone levels [1] and to the expression of *PSCK 2* in older barn owl nestlings [95, 99]. Finally, we cannot exclude that the expression of *GR* and *MR* are independent of the expression of *PCSK 2* and that the relation between those genes may occur through a pathway that is independent of the melanocortin system.

Alternatively, the link between melanin-based traits and the expression of *MR* and *GR* could arise because the genes regulating the deposition of melanin pigments and these receptors are co-expressed, owing for instance to a common factor regulating the expression levels of melanogenic and the receptors binding glucocorticoids. In this case, we would have expected a negative association between the expression levels of melanogenic-related genes and *MR* and *GR* genes, given that the expression of the latter genes in this study and in a previous study in house sparrows is smaller in more melanised individuals (Figure 3). In our study, however, we did find significant associations between glucocorticoids receptors (*MR, GR*) and genes involved in the production of melanin pigments (*TYR, OCA2* (PC 1), *MITF-M* (PC 2), Figure 1 & 2) and in the biogenesis and development of melanosomes (*MLANA, PMEL* (PC 2), Figure 1 & 2), genes coding for proteins involved in the early steps of synthesis, storage and transport of melanin pigments [110]. However, such associations were positive rather than negative, what we would have expected given the negative association between spot diameter and glucocorticoid receptors (*MR, GR*). This finding suggests that the covariation between melanin-based traits and the expression of *MR* and *GR* is not mediated by the melanogenic-related genes considered here.

The absence of evidence for a negative relationship between *MR* and *GR* with the expression of melanogenic-related genes could be due to the timing of feather sampling. Although we tried to pinpoint the time when individuals produce the black spots, we do not have the certitude that all individuals were producing the spots at the time when we plucked feathers. This is supported by the fact that we found little evidence for a positive relationship between spot size and PC 1 (β= 0.01, 95% CrI [-0.06; 0.08]) and PC 2 (β= −0.04, 95% CrI [-0.13; 0.05]), PC1 being the principal component the most representative of the expression of melanogenic genes (see Table 1). This suggests that the expression levels of melanogenic genes at the time when we collected the feathers is not yet entirely representative of the final colouration that the individuals will display. Future studies accounting for temporal evolution of gene expression in feathers may help to better understand why we observed an association between the expression of *GR* and *MR* measured at an early age and melanin-based traits but not a similar association with the genes associated to the production of melanin pigments.

Although the potential for corticosterone to affect the expression of ornaments, such as melanin-based traits has been demonstrated in different studies [47, 58, 88] and taxa [40, 73], we do not know how corticosterone directly or indirectly affects the deposition of melanin pigments, including the role of MR and GR in this process. Assuming that a lower expression of *GR* and *MR* in the tissues corresponds to fewer receptors available for glucocorticoids, large-spotted individuals could reduce the long-term negative effect of high corticosterone levels [47, 58, 88]. However, the action of glucocorticoids (corticosterone in birds) in the skin or feather tissues may be limited or exist only when levels are elevated given that, in contrast to brain tissues where glucocorticoids can bind to MR and GR, the full occupancy of GR and MR by glucocorticoids in the skin is prevented by the action of 11ß-hydroxysteroid dehydrogenase type 2 (11ß-HSD2) that converts corticosterone to inert intracellular products of corticosterone [18, 48]. Future studies should investigate to what extent corticosterone effects, at the integumentary level are actually mediated by GR and MR receptors in order to clarify how corticosterone could be affecting colouration at the level of the integument *via* binding to its receptors. Interestingly, in humans, 11ß-HSD2 limits also the access of GR and MR to aldosterone, which regulates salt homeostasis, epidermal and hair development [32, 34, 44, 75]. This opens an alternative pathway for GR and MR receptors to affect melanin-based coloration via altering the development of the integument where melanin is deposited. GR and MR act as transcription factors that once bound to their ligands (aldosterone, corticosterone) are critical for the development of hair and skin cells. For instance, overexpression of *MR* and *GR* in keratinocyte of mice can induce skin atrophy resulting in the thinning of the upper layers of the skin causing them to be more fragile [76, 93]. Although in contrast to humans and mice [32, 75] the roles of GR and MR in the development of feathers are unknown, variation in the expression of glucocorticoid and mineralocorticoid receptors in tissues of growing feathers might also have some effects on the development of feathers for instance in the rate of feather growth [70]. These effects could induce structural differences in the feathers [91] leading to differential expression of melanin-based traits, an hypothesis that could be addressed in future studies.

From a wider perspective, variability in the availability of corticosterone receptors in other tissues regulated by corticosterone may also have important physiological consequences. The availability of *GR* and *MR* in the hypothalamus plays a crucial role in the sensitivity of HPA axis as it determines the capacity of the system to come back to normal corticosterone basal levels [25] and to avoid the damaging effects of high and prolonged corticosterone levels. Individuals with lower *GR* and *MR* expression are assumed to have fewer available receptors and thus, have a reduced feedback signal when facing acute stress. A mitigated negative-feedback response implies that individuals will be exposed to higher concentration of corticosterone, which could have serious consequences on their health [7, 26, 65]. Differences in the expression levels of corticosterone receptors (*GR, MR*) and corticosterone plasma levels during an acute stress between differently melanized individuals could indicate that barn owls have different strategies to cope with stress, which could be signalled by colour traits [56]. Large-spotted barn owls may have higher peaks of corticosterone during an acute stress response but express glucocorticoid receptors at higher levels which enables them to recover from an acute stress faster than small-spotted individuals who, on the other hand, may produce less corticosterone during an acute stress. Although GR and MR are, to some extent, similarly expressed in different tissues throughout the body [57], additional studies linking *GR* and *MR* expression levels in tissues involved in the regulation of the HPA axis, including metabolic (*i.e.,* liver, kidney, subcutaneous fat and muscles) and brain tissues, to melanin-based traits and melanocortin related genes will help us to better understand how differently melanized individuals modulate the HPA axis and how melanin-based coloration evolve as stress-related signals.

## Acknowledgement

We would like to thank Robin Séchaud and the numerous field assistants for their help on the field and to Hannes Richter for his help and guidance in the laboratory analysis. Funding was provided by Swiss National Science Foundation (grant n° 31003A-120517 to AR). The study was performed under legal authorization of the ‘Service vétérinaire du canton de Vaud’.

## Author Contributions

A.R. obtained funding. A.R.; P.B. and A.-L.D. conceived and designed the study; P.B. conducted fieldwork; P.B., A.-L.D. and C.S. conducted all the genetic analysis; P.B. conducted the statistical analyses; P.B and A.R. wrote the manuscript with important contributions of L.M.S.-J. and A.-L.D. All authors read and provided input on the manuscript.

